# GhostFold: Accurate protein structure prediction using structure-constrained synthetic coevolutionary signals

**DOI:** 10.1101/2025.10.13.682177

**Authors:** Nitesh Mishra, Bryan Briney

## Abstract

The accuracy of protein structure prediction models such as AlphaFold2 is tightly coupled to the depth and quality of multiple sequence alignments (**MSAs**), posing a persistent challenge for proteins with few or no identifiable homologs. We present GhostFold, a method for conjuring structure-constrained synthetic MSAs from a single amino acid sequence, bypassing the need for traditional homology searches. Leveraging the ProstT5 protein language model and the 3Di structural alphabet, GhostFold projects a query sequence into a tokenized structural representation and iteratively back-translates to generate an ensemble of diverse, fold-consistent sequences. These synthetic alignments (**pseudoMSAs**) encode emergent coevolutionary constraints that are sufficient for high-accuracy structure prediction of difficult targets such as orphan proteins and hypervariable antibody loops. GhostFold consistently matches or exceeds the performance of MSA-based and language model-based structure predictors while being computationally lightweight and independent of large sequence databases. Notably, we observe a decoupling of confidence metrics (e.g., pLDDT) from prediction accuracy when using pseudoMSAs, suggesting that AlphaFold2’s internal confidence calibration is strongly influenced by the statistical properties of natural sequence alignments. These results establish that structure-guided synthetic MSAs can functionally substitute for evolutionary data, offering a scalable and generalizable solution to one of the central limitations in computational structural biology. GhostFold represents a shift from passive data mining to intelligent sequence synthesis, redefining how structural priors are encoded in deep learning-based protein folding.

## INTRODUCTION

The advent of deep learning-based protein structure prediction, exemplified by AlphaFold2 (**AF2**) and its derivatives, has reshaped the field of structural biology [1]. Central to the success of these models is the input multiple sequence alignment (MSA), which encodes coevolutionary constraints critical for inferring accurate structures. Accordingly, much effort has focused on improving MSA construction pipelines through iterative profile searches, metagenomic mining, and database fusion and methods such as DeepMSA2, D-I-TASSER, and others have enabled deeper alignments by searching larger and more diverse datasets [2,3]. However, such strategies face practical limitations; increasing database size yields diminishing returns for sequences with few or no detectable homologs and demands ever larger storage and compute resources.

Parallel efforts have explored using the embeddings produced by protein language models (**pLMs**) to predict structure directly from single sequences. Models such as ESMFold [4], OmegaFold [5], and RGN2 [6] avoid MSA generation entirely and offer rapid inference, but are typically less accurate than MSA-based methods like AF2. Their limitations have been particularly evident in recent CASP experiments, where pLM-based predictors consistently lagged behind MSA-dependent methods [7,8]. Fundamentally, pLM-based models operate without explicit representations of residue-residue coevolution, limiting their effectiveness for sequences where evolutionary constraints are the primary source of structural information. Domain-specific predictors can mitigate these shortcomings in specialized settings like antibodies, but at the cost of generalizability [9,10].

The reliance on ever-expanding sequence databases as a primary lever for improving prediction accuracy has become increasingly untenable. Petabase-scale datasets have been integrated into search pipelines [11], yet structure prediction performance for orphan proteins and complex sequences remains stubbornly low. This suggests an upper bound on the information content that can be mined from natural sequence space and underscores the need for alternatives that do not depend on historical evolutionary data. Previous works, such as MSA-Generator [12], MSAGPT [13], and PLAME [14], have attempted to address this challenge by generating virtual MSAs using an encoder-decoder framework. While these models can synthesize plausible sequences that enhance the diversity of an existing MSA, they still depend on a homology-derived alignment as a starting point. In other words, these approaches augment rather than replace the need for conventional MSAs, limiting their applicability for true orphan or synthetic protein sequences without natural homologs. A framework that removes this dependency altogether, generating structurally informative alignments directly from a single sequence, remains a key unmet need.

Here, we present GhostFold, a structure prediction framework that relies on structure-informed pseudoMSAs produced *de novo* from a single query sequence. Our approach leverages ProstT5 [15], a bilingual pLM trained to translate between amino acid sequences and discrete structural representations. The FoldSeek 3Di structural alphabet [16] encodes tertiary residue interactions, enabling projection of a sequence into a tokenized representation of its structure. From this tokenized representation, novel, fold-consistent amino acid sequences are sampled and assembled into a pseudoMSA. This process recapitulates the types of coevolutionary constraints typically inferred from natural alignments, while remaining agnostic to the number and diversity of natural sequence homologs.

We show pseudoMSAs generated by GhostFold consistently improve the accuracy of predictions for orphan and low-homology proteins. These improvements do not arise from increased alignment depth per se, but from the sharpening of the coevolutionary signal introduced by the generative process. GhostFold performs extremely well across a broad spectrum of prediction challenges: in cases where pLM-based models produce the best results, GhostFold performs equally well; in settings where MSA-dependent models excel, GhostFold matches or exceeds their performance. This lack of trade-off underscores that the improvements are not confined to edge cases or niche applications. GhostFold either improves prediction accuracy or performs on par with existing methods, with no observed regressions. Importantly, this framework provides a scalable and computationally efficient alternative to database-driven pipelines, requiring no homology search and minimal input preprocessing. We further argue that continued reliance on database expansion is unsustainable and increasingly ineffective for the most challenging prediction problems. Structure-aware synthetic data offers a practical path forward, complementing architectural advances in prediction models with improved control over input quality and diversity.

## RESULTS

### Zero-shot generation of structurally coherent pseudoMSAs

Our methodology is centered on a zero-shot framework that generates a high-quality pseudoMSA from a single query sequence, circumventing the need for computationally expensive and sometimes fruitless homology searches over massive sequence databases. We illustrate this pipeline using the orphan protein ToPI1 inhibitor [17] (PDB ID: 6MRQ), a 33-residue β-grasp fold (***Fig 1a-b***). In the initial stage of pseudoMSA construction (AA to 3Di), our goal is a thorough exploration of the structural representation space centered around the most probable structural scaffold. Simply generating the top *N* most likely 3Di representations would yield a set of highly similar tokenized structures, providing limited information for the final alignment. To build a rich pseudoMSA, we instead need to sample a diverse set of high-probability structural projections that are all compatible with the input sequence’s local and global fold. To achieve this, we require a search algorithm that can discover multiple high-quality, diverse outputs from a large number of 3Di projections of a single input sequence. Standard beam search maintains a predetermined number of the best options, or “beams,” at each step, preventing the model from committing to a single suboptimal path too early. A purely deterministic beam search protocol would still converge on a limited set of very similar, high-probability structures, failing to provide the desired diversity. Instead, we utilized beam search with multinomial sampling [18] where beams are ranked by their length-normalized sum of log probabilities. This hybrid approach expands on the vanilla beam search framework by introducing stochasticity at each generation step. This ensures that our exploration of the 3Di projection space does not collapse into repetitive outputs but instead produces a varied ensemble of high-quality structural representations. In the second state of pseudoMSA construction (3Di to AA), our objective shifts from exploration to high-fidelity sequence reconstruction, so we employ a direct multinomial sampling strategy. By sampling from the model’s predicted distribution at each step, this approach generates sequences that are more statistically aligned with natural proteins. This deliberate shift from a broad, beam-based exploration to distributional sampling for the amino acid reconstruction ensures the creation of novel proteins that both thoroughly sample the conformational space compatible with the input sequence’s fold and coherently translated in sequence.

**Figure 1.**
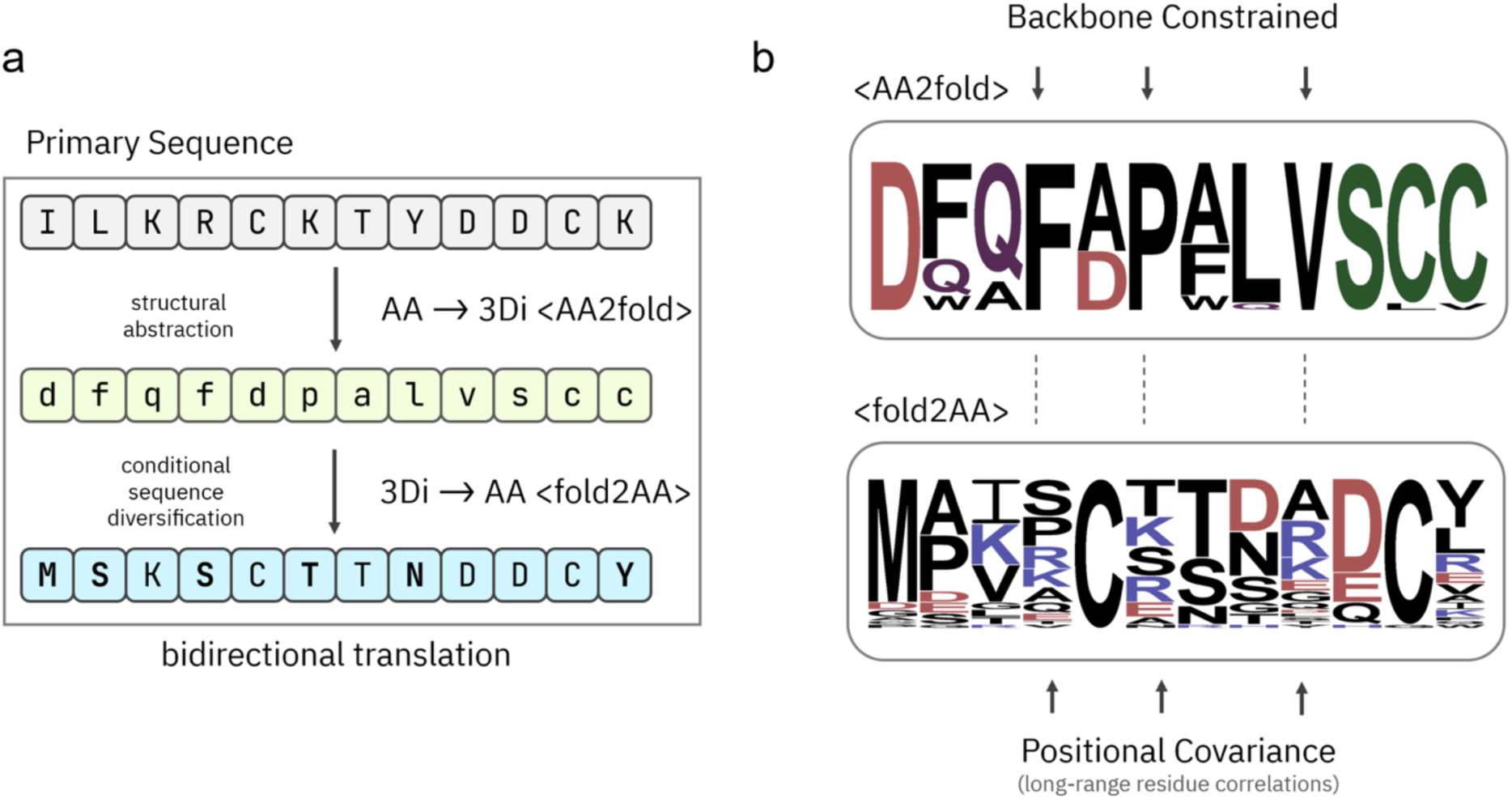
Bidirectional translation between sequence and structure enables zero-shot generation of structurally coherent pseudoMSAs. (a) Schematic overview of the hierarchical generation process used to construct pseudoMSAs from a single input sequence. Beginning with a primary amino acid sequence (top, grey), the model first performs structural abstraction by translating into a structural latent representation using the 3Di alphabet (middle, green) via the AA to 3Di process (<AA2FOLD>). This abstraction captures fold-compatible structural features while allowing for backbone-constrained variation. Next, conditional sequence diversification is performed by projecting from 3Di back to amino acids (bottom, blue) via the 3Di to AA process (<FOLD2AA=), enabling diverse yet fold-consistent synthetic sequences. This bidirectional translation allows sampling of multiple high-quality, structurally coherent sequences from a single query. (b) Information content of the latent and reconstructed sequence spaces visualized as sequence logos. The upper logo (<AA2FOLD>) depicts the distribution of 3Di-derived structural abstractions constrained by the input sequence’s backbone. The lower logo (<FOLD2AA=) shows the resulting sequence distribution after back-translation to amino acids, which reflects both local backbone compatibility and positional covariance, capturing long-range residue-residue correlations. Together, these components illustrate how structural coherence and evolutionary plausibility are maintained throughout the pseudoMSA generation pipeline.

To further enrich the diversity and information content of the final pseudoMSA, we augmented this core generative loop with several additional strategies. Our pipeline includes iterative stochastic decoding across a spectrum of sequence coverages, ranging from 40% to 100%, to generate both full-length sequences and sequence fragments that capture local structural motifs and coevolutionary signals (***Fig S1a–c***). Furthermore, we introduce controlled evolutionary drift through targeted mutagenesis, using established substitution matrices like BLOSUM62, PAM250 and MEGABLAST to generate evolutionarily plausible sequence variants. The aggregated set of generated sequences are filtered to reduce redundancy and distill a final, high-quality pseudoMSA (***Fig S1b–d***). The sequence identity of pseudoMSA sequences to the original query remains in the 30-50% range, indicating that the generated diversity is well-constrained around the input fold (***Fig S1c***). This controlled diversity is further detailed in the sequence logo of the pseudoMSA (***Fig S1d***), which reveals distinct amino acid preferences at each position, reflecting the learned physiochemical constraints of the β-grasp fold. For instance, conserved glycine and proline residues, critical for turns, are clearly represented in the pseudoMSA.

Crucially, this synthetically generated alignment contains a strong, emergent co-evolutionary signal, which is the essential ingredient for accurate structure prediction. While the contact maps derived from individual back-translated sequences are sparse and noisy (***Fig S1a***), their aggregation and filtering reveal a clear, structured co-evolutionary map (***Fig S1b***). This map exhibits strong off-diagonal signals corresponding to contacts between distal residues that form the core of the fold. The emergence of this coherent signal from a diverse but structurally constrained sequence ensemble provides the necessary information for downstream modelling with AlphaFold2. This comprehensive approach allows for the exploration of a vast ensemble of sequences that are thermodynamically compatible with a given fold, creating informative surrogates that empower downstream structure prediction.

### Benchmarking against state-of-the-art single-sequence structure predictors

To validate the effectiveness of our pseudoMSA generation pipeline, we benchmarked its performance on the Orphan benchmarking dataset (see ***Methods***), a challenging set of 36 naturally occurring proteins that do not have sequence homologs [5,19]. We created GhostFold by providing pseudoMSA inputs to AF2 using the community led ColabFold framework [20]. GhostFold was then benchmarked against AF2 (in both single-sequence and full MSA mode) and two leading single-sequence structure prediction models, ESMFold, and OmegaFold, which use protein language model embeddings instead of MSAs to extract co-evolutionary signals [4,5].

The results demonstrate a marked improvement in prediction accuracy when using our synthetically generated pseudoMSAs. GhostFold consistently produces predictions with lower root mean square deviation (RMSD) from the ground-truth structure, indicating higher predictive accuracy (***Fig 2a*** and ***Fig S2a–b***). While all methods show a wide performance range, GhostFold’s predictions are more concentrated in the lower RMSD tiers compared to the other models (***Fig S2a***). Of note, for 12 of 36 proteins, none of the methods predicted accurate structure though GhostFold got majority of the domains for several of the failed cases (Fig S3).

**Figure 2.**
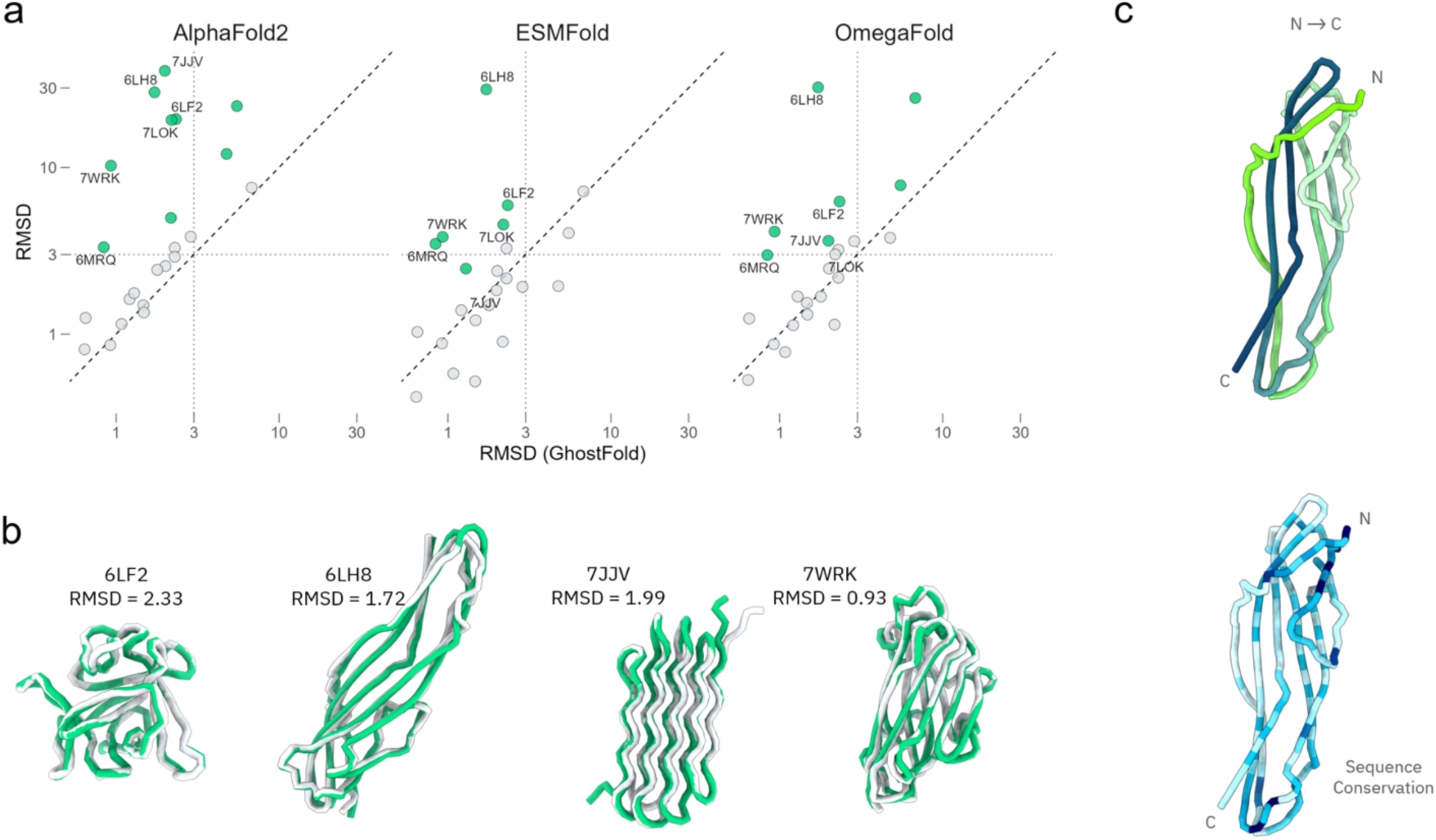
GhostFold outperforms single-sequence predictors on orphan proteins. (a) RMSD comparison between GhostFold and three leading single-sequence prediction methods: AlphaFold2 (AF2, single-sequence mode), ESMFold, and OmegaFold. Each panel plots the RMSD of GhostFold predictions (x-axis) against the indicated method (y-axis). The dashed diagonal represents equal performance; points above the line indicate superior prediction accuracy by GhostFold. A dotted threshold line at 3 Å RMSD demarcates high-confidence models. Green-highlighted points correspond to examples where GhostFold predictions achieved RMSD < 3 Å while the comparator method failed (RMSD > 3 Å). These examples are labeled by PDB ID. Gray points indicate comparable performance. (b) Structural overlays for four representative proteins where GhostFold achieved high-accuracy predictions. Native structures are shown in gray, and GhostFold predictions are in green. The corresponding PDB IDs and RMSD values are indicated below each model. Examples include complex beta-rich topologies such as 6LF2, 6LH8, 7WRK, and amyloid protein 7JJV. (c) Sequence conservation mapping of the pseudoMSA used by GhostFold, illustrated for the 6LH8 example. Left: predicted structure (light to dark green gradient denotes N- to C-terminal directionality). Right: same model colored by pseudoMSA conservation (blue gradient), where darker blue indicates higher sequence conservation. Despite limited conservation (∼50% at most sites), GhostFold successfully recovers the native fold.

For AlphaFold2, ESMFold and OmegaFold, majority of data points fall above the diagonal line, highlighting higher RMSD values for the indicated method, indicating a superior prediction by GhostFold for most targets in the dataset (***Fig 2a***). This systematic improvement is particularly notable against the standard AlphaFold. Furthermore, GhostFold achieves a significantly lower RMSD (in green) for the majority of the orphan proteins compared to AlphaFold2, ESMFold and OmegaFold (***Fig 2b***). This performance advantage is highlighted by specific examples (***Fig 2c***). For the more complex beta-sandwich folds of 6LF2, 6LH8, and 7WRK, our method yields a model with an RMSD of <2 Å, successfully capturing the intricate topology where other methods might struggle. Even for challenging targets like the amyloid-like protein 7JJV, composed of tightly packed beta-sheets, GhostFold produces a commendable model with an RMSD of 1.99 Å, correctly predicting the overall fold and strand arrangement. Of note, for the majority of the successful predictions, the pseudoMSA showed 50% sequence conservation at most sites with ***Fig 2d*** highlighting the case of 6LH8 where pLM structure prediction models, ESMFold and OmegaFold, failed drastically.

An important aspect of evaluating deep learning models is assessing their internal confidence metrics. We investigated the relationship between the final model accuracy (RMSD) and the two standard AlphaFold2 confidence scores: the per-residue confidence (pLDDT) and the predicted TM-score (pTM).

Our analysis revealed a striking difference in how model confidence relates to accuracy when using natural versus synthetic MSAs. For standard AlphaFold2, which uses homology-searched MSAs, there is a strong correlation between higher pLDDT scores and lower RMSD values (R² = 0.80), confirming that the pLDDT is a reliable estimator of model accuracy (***Fig S2c***). However, when using GhostFold pseudoMSAs, this correlation weakens (R² = 0.44). High-accuracy models (low RMSD) do not consistently receive high pLDDT scores, and several predictions assigned high pLDDT scores are, in fact, inaccurate. The pTM score, which assesses the global fold, was much less affected by GhostFold’s pseudoMSAs (***Fig S2d***). Collectively, these results show that by providing a structurally consistent and diverse pseudoMSA, GhostFold effectively guides the prediction model towards the native state, overcoming the limitations of single-sequence approaches and shallow MSAs for orphan proteins.

### GhostFold excels without iterative refinement

To further assess GhostFold performance, we used the Design55 dataset [19], a collection of 55 human-designed proteins that lack natural homologs and thus serve as an ideal testbed for single-sequence prediction capabilities. Unlike naturally occurring orphan proteins, these sequences have often been optimized for exceptional stability, posing a unique challenge. Our analysis reveals that providing a pseudoMSA not only improves final accuracy but also fundamentally alters the prediction dynamics within AlphaFold2, rendering its iterative refinement process largely redundant.

A key component of the AlphaFold2 architecture, particularly in the absence of a deep MSA, is its recycling feature, where the model’s output is fed back as input to iteratively refine the structure. For the Design55 dataset, AlphaFold2 in single sequence mode shows a heavy reliance on this process (***Fig 3a–b***). The model’s confidence metrics, pLDDT and pTM, steadily increase over several recycling iterations, indicating iterative convergence towards a final predicted structure. In contrast, GhostFold arrives at its optimal prediction almost immediately. When provided with a synthetic pseudoMSA, the pLDDT and pTM scores start at a high level from the first cycle (recycle 0) and remain essentially flat throughout the recycling process (***Fig 3a–b***). This indicates that the pseudoMSA provides the Evoformer module with a rich and coherent structural signal from the outset, reducing the need for iterative refinement. This pattern is universal across all five AF2 model variants (***Fig S4a–b***).

**Figure 3.**
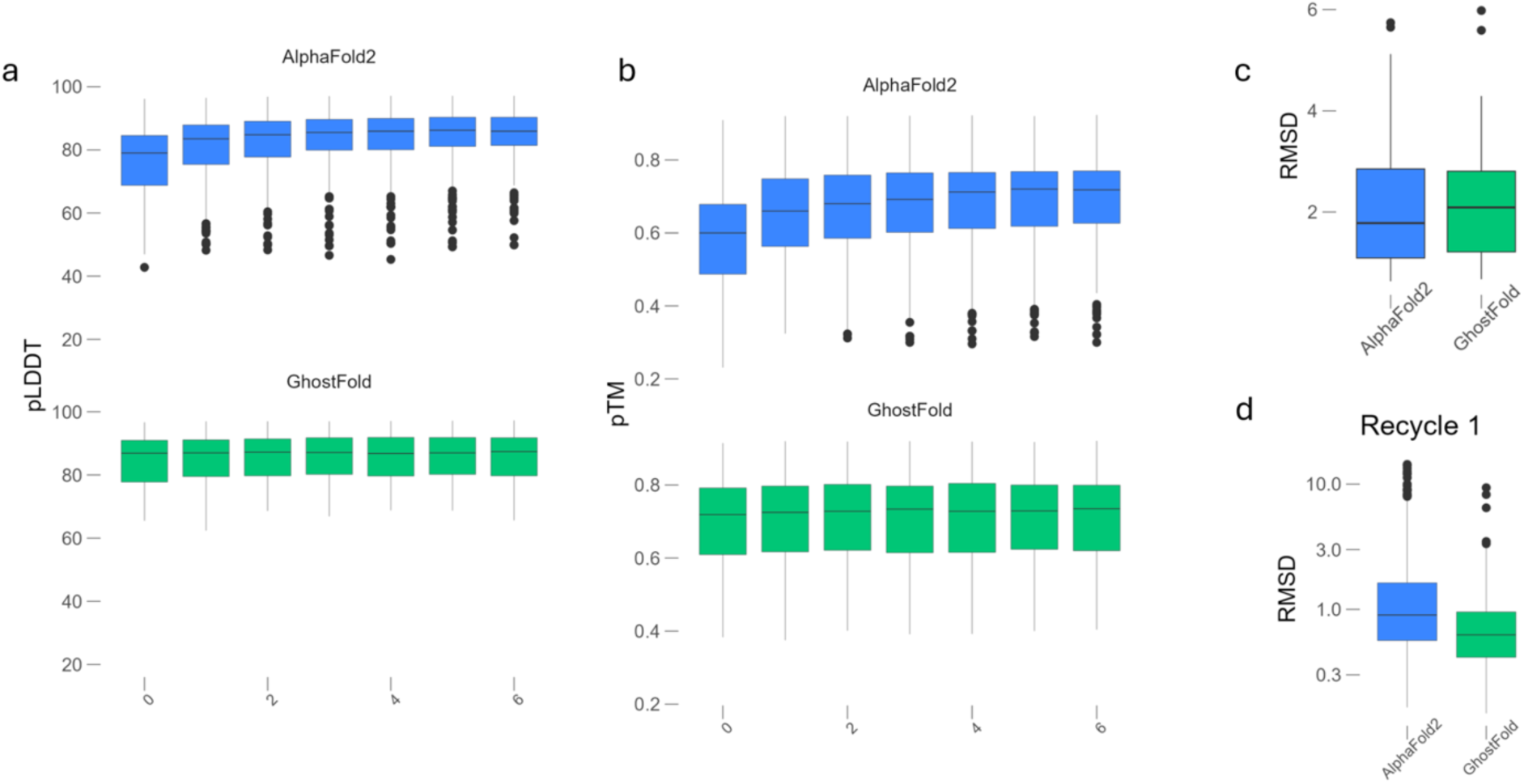
GhostFold eliminates the need for iterative refinement on de novo designed proteins. (a) Per-residue confidence (pLDDT) scores across AlphaFold2 (top, blue) and GhostFold (bottom, green) for the Design55 dataset, plotted over recycling iterations (0 to 6). AlphaFold2 predictions show a gradual increase in pLDDT with recycling, indicative of progressive model refinement. In contrast, GhostFold begins with high pLDDT scores from recycle 0, which remain stable throughout subsequent recycles, suggesting that the pseudoMSA provides sufficient structural signal from the outset. (b) Predicted TM-score (pTM) values over the same recycle steps, showing a similar trend. AlphaFold2 exhibits a steady increase in global fold confidence with each recycle, whereas GhostFold achieves high pTM immediately with minimal change thereafter. (c) Final model accuracy comparison using RMSD for the best-ranked structure after full recycling. GhostFold and AlphaFold2 produce comparable median RMSDs, though GhostFold shows slightly tighter distribution. This demonstrates GhostFold’s ability to match or exceed the performance of single-sequence AlphaFold2, even without depending on iterative recycling. (d) RMSD comparison after the first recycling step (Recycle 1) shows a striking difference: GhostFold achieves near-final accuracy (median RMSD ∼1.0 Å), while AlphaFold2 lags significantly, with a median RMSD closer to 3 Å and greater variance. This supports the conclusion that the iterative refinement in AlphaFold2 is compensating for insufficient input signal, a limitation that GhostFold overcomes via pseudoMSA augmentation.

This high initial confidence translates directly to accurate models as evident by comparable RMSDs for GhostFold and AlphaFold2 in single sequence mode (***Fig 3c***). Interestingly, at the first recycle step, GhostFold models already exhibit a tight distribution of low root-mean-square deviation (RMSD) values, with a median RMSD around 1.0 Å. Standard AlphaFold2, even after recycling, lags significantly, producing models with a median RMSD nearly three times higher and a much wider variance (***Fig 3d***). This confirms that the iterative refinement in the single-sequence modality is a compensatory mechanism for a weak initial structural signal, a limitation that GhostFold effectively overcomes. While prior work correctly identified that recycling was key to AlphaFold2’s performance on this dataset, our findings demonstrate that a high-quality synthetic MSA is an effective alternative, providing the necessary structural information upfront for a direct and accurate prediction.

### Assessing GhostFold performance using an oracle-guided benchmark

To better understand the performance limits of structure prediction from synthetic data, we designed an “oracle” experiment using ProteinMPNN [21], a state-of-the-art model that generates amino acid sequences conditioned on a fixed 3D backbone structure. By providing ProteinMPNN with ground-truth experimental structures from the Orphan dataset, we generated optimal pseudoMSAs that, to the extent ProteinMPNN is capable, provide the correct structural information. This allowed us to ask a critical question: how does AlphaFold2 perform when given an ideal coevolutionary signal?

As expected, the coevolution maps derived from ProteinMPNN-generated MSAs are exceptionally clean. For challenging targets like 7JJV and 6LH8, where GhostFold pseudoMSAs produce a usable but noisy signal, the ProteinMPNN-based MSA yields a sharp, nearly perfect representation of the native contact map (***Fig 4a–b***). This represents a theoretical upper bound on the quality of the input signal. The final prediction accuracy, however, reveals a surprising paradox. Despite being fed this near-perfect input, the ProteinMPNN-augmented AF2 pipeline still failed to produce an accurate model for approximately 20% of the targets in the orphan set (***Fig 4c–d***, “AF2 rcsb”). This suggests that for a subset of difficult proteins, providing a perfect coevolutionary map is not sufficient to guarantee a correct final structure, highlighting potential limitations within the AF2 framework itself for accurately predicting certain complex topologies.

**Figure 4.**
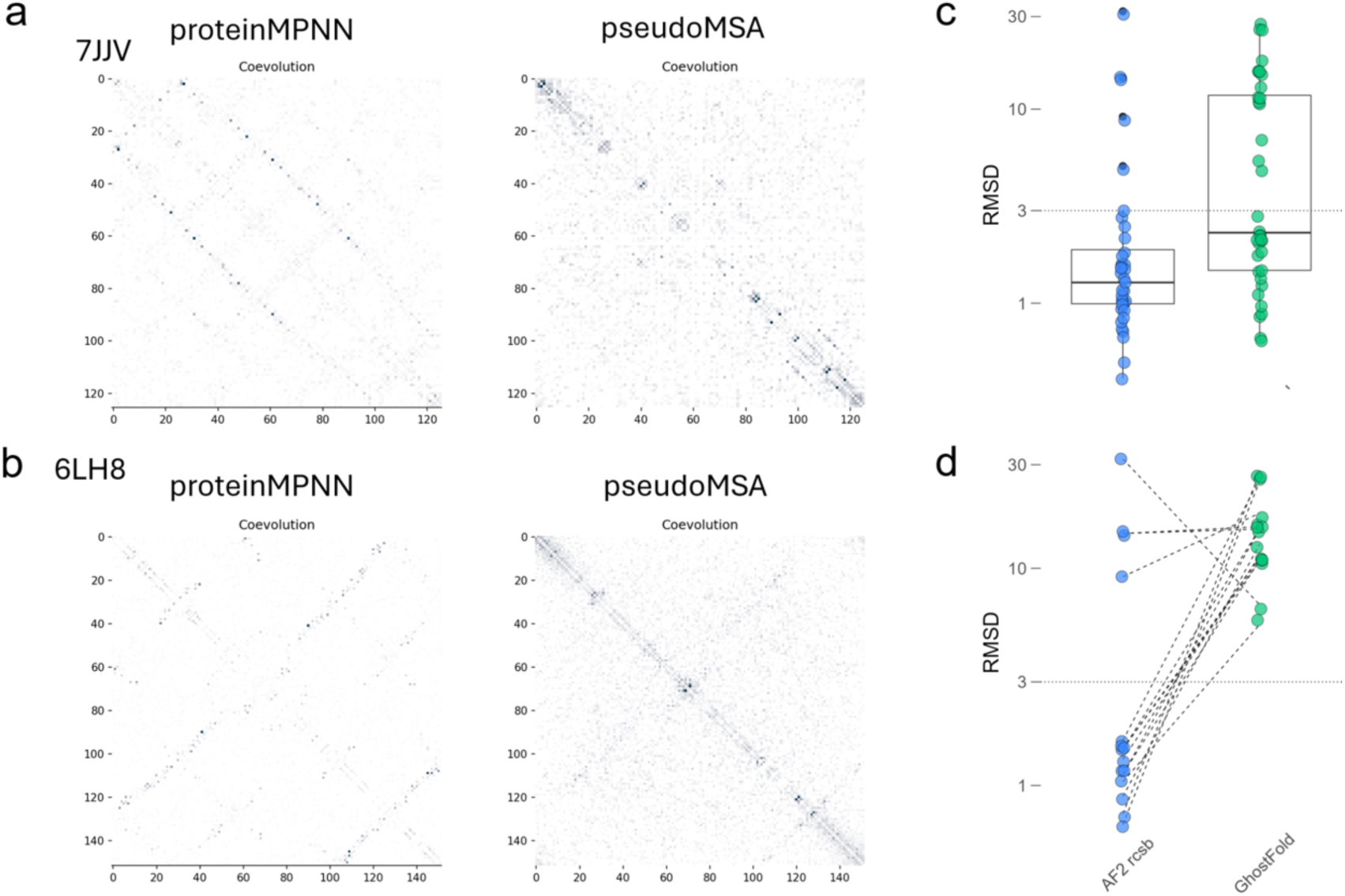
GhostFold approaches the theoretical upper bound of structure prediction from synthetic coevolutionary signals. (a–b) Comparison of coevolution signal derived from pseudoMSAs generated via GhostFold (right) and an oracle-based method using ProteinMPNN (left), for two representative orphan proteins: 7JJV (a) and 6LH8 (b). For the ProteinMPNN approach, sequences were sampled using the ground-truth PDB structure as input, producing MSAs that encode an idealized representation of the native topology. The resulting coevolution maps show sharp and accurate contact patterns. GhostFold pseudoMSAs, though noisier, still recover key structural contacts. Axes represent residue–residue positions; pixel intensity reflects coevolutionary coupling strength with darker colors implying strong coevolutionary signal. (c) RMSD comparison between AlphaFold2 predictions using MSAs generated by ProteinMPNN (“AF2 rcsb”, blue) and GhostFold (green) across the orphan protein set. Boxplots summarize the RMSD distribution for each method, with individual protein points overlaid. Although ProteinMPNN-based inputs yield lower median RMSD overall, GhostFold’s performance is comparable for a large subset of targets despite lacking ground-truth structural input. (d) Per-target RMSD comparison for cases where GhostFold failed (RMSD > 5 Å). Even when provided with the ideal pseudoMSA derived from the native structure (via ProteinMPNN), AlphaFold2 still fails to accurately predict ∼20% of these structures, suggesting that architectural limitations—rather than MSA quality—may underlie persistent errors in structural prediction for certain challenging folds. Dotted lines connect RMSDs for the same protein across both input conditions.

This finding places the performance of GhostFold into a new perspective. Our method, which operates with no prior knowledge of the ground-truth structure and generates a comparatively “fuzzy” structural signal, performs only marginally worse compared to ideal pseudoMSAs generated using structural information, failing on just ∼10% more targets than the oracle-guided approach. The fact that GhostFold’s performance is so close to this theoretical benchmark underscores the remarkable effectiveness of its structure-sampling strategy. It implies that the specific combination of sequence diversity and structural information generated by ProstT5 tokenization and back-translation may be particularly well-suited for guiding AF2 predictions, and that simply maximizing the cleanliness of the coevolutionary signal is not the ultimate determinant of success.

### GhostFold accurately predicts hypervariable antibody CDRH3 loops

Antibody structure prediction represents a uniquely challenging frontier for protein modeling. The primary difficulty lies in the hypervariable complementarity-determining regions (CDRs), particularly the heavy chain CDR3 (CDRH3) loop, which is often long, structurally diverse, and lacks a canonical conformation. Due to its central role in antigen recognition, accurately modeling the CDRH3 loop is paramount. However, its hypervariability means it often has no evolutionary homologs outside of closely related antibodies, rendering it analogous to an orphan peptide embedded within a stable scaffold. This makes its prediction notoriously challenging, often requiring massive computational sampling to achieve accuracy, a limitation acknowledged even in state-of-the-art models like AlphaFold3 [22,23]. We hypothesized that GhostFold’s ability to generate structurally plausible sequences for orphan regions could help overcome this long-standing computational hurdle.

To test this, we curated a challenging benchmark set of 71 non-redundant nanobody structures deposited in the PDB during 2025. We compared the performance of GhostFold against a standard AlphaFold2 pipeline using MMseqs2-generated MSAs [20,24,25]. The results demonstrate that GhostFold not only improves accuracy but, more importantly, achieves a high level of consistency in its predictions.

A striking example is the case of 9G9M (chain C), a nanobody with a long and complex 18-residue CDRH3 loop (***Fig 5a***). GhostFold predicts the entire nanobody with a global RMSD of 0.68 Å. Critically, the challenging CDRH3 loop itself is modeled with remarkable precision, achieving an RMSD of just 0.41 Å from the crystal structure (***Fig 5a***, left). The true power of our approach is revealed by examining the ensemble of predicted models. For GhostFold, the five models generated by AlphaFold2 are virtually indistinguishable from one another, appearing as a single ribbon in the structural overlay and yielding an ensemble CDRH3 RMSD of 0.53 Å (***Fig 5a***, right). This indicates that the pseudoMSA provides such an unambiguous structural signal that the model arrives at the correct solution practically deterministically.

**Figure 5.**
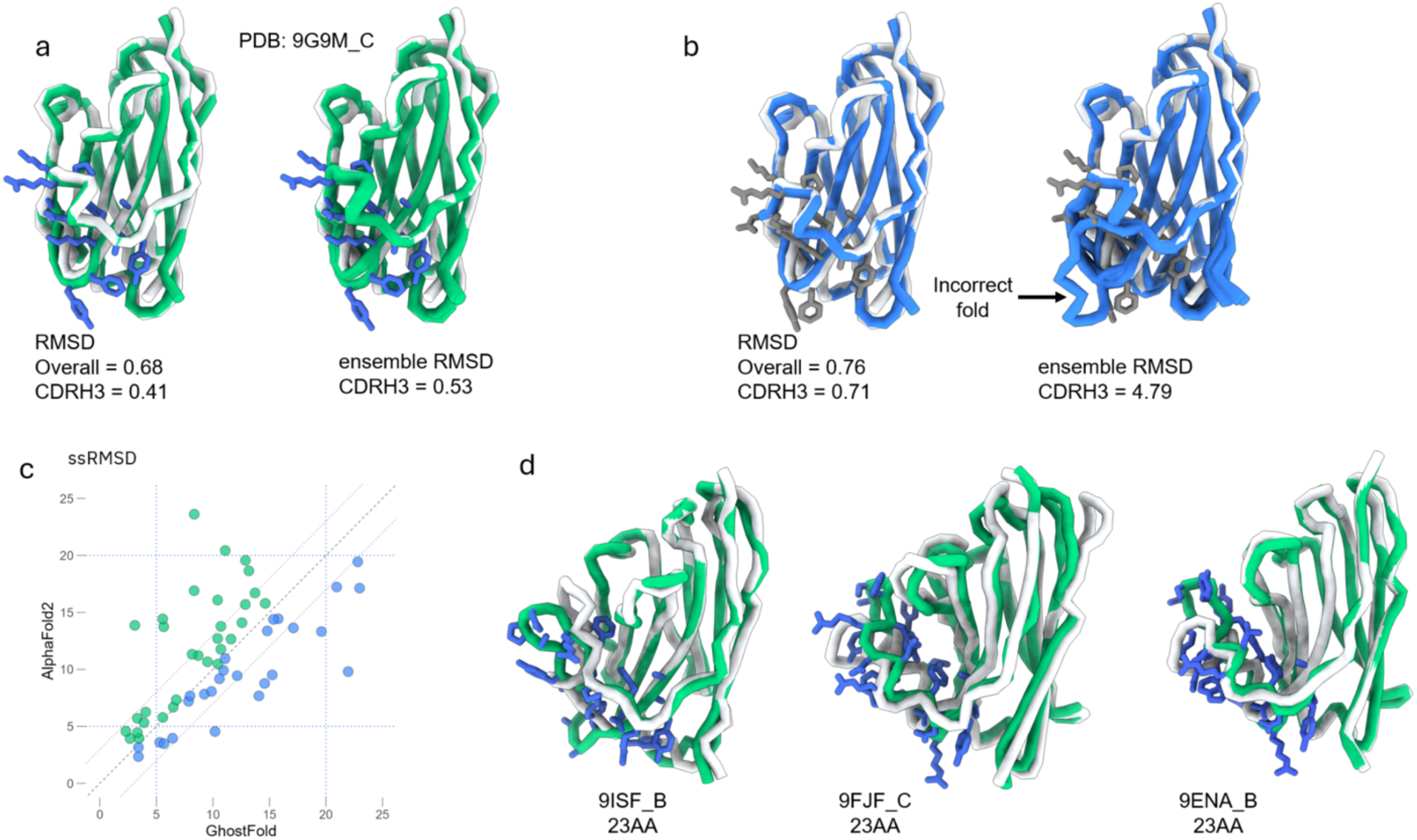
GhostFold enables accurate and ensemble-consistent prediction of hypervariable CDRH3 loops. (a) Representative example of GhostFold performance on nanobody 9G9M_C. Left: Overlay of predicted structure (green) and crystal structure (gray), with the residues in CDRH3 loop highlighted in blue. GhostFold achieves high global accuracy (RMSD = 0.68 Å) and precise modeling of the CDRH3 loop (RMSD = 0.41 Å). Right: Structural ensemble across the top five GhostFold models shows minimal deviation (ensemble CDRH3 RMSD = 0.53 Å), reflecting high consistency enabled by the pseudoMSA. (b) Corresponding result from the standard AlphaFold2 pipeline using MMseqs2-derived MSAs. While the overall RMSD is comparable (0.76 Å), the CDRH3 loop deviates more (RMSD = 0.71 Å) and the ensemble prediction is highly variable (ensemble CDRH3 RMSD = 4.79 Å). The five models diverge substantially in the loop region, failing to converge on the correct conformation. Predicted models in blue, CDRH3 residues in gray. (c) Per-target comparison of ensemble consistency between GhostFold and AlphaFold2 across 71 nanobodies. Each point represents a nanobody, with the x- and y-axes showing the summed squared RMSDs (ssRMSD) for GhostFold and AlphaFold2, respectively. Cases where Ghostfold had better ensembles are colored green while cases where AlphaFold2 with mmseqs was better are shown in blue. (d) Structural overlays of three nanobody targets with long CDRH3 loops (23 amino acids): 9ISF_B, 9FJF_C, and 9ENA_B. In each case, GhostFold predictions (green) match closely to the native structure (gray) with accurate modeling of the long CDRH3 loops (residues in blue). The minor deviations occur predominantly at flexible loop tips.

In contrast, the standard AlphaFold2-MMseqs2 pipeline produces a model with a comparable global RMSD (0.76 Å) but fails on the tip of the CDRH3 loop, which deviates by 0.71 Å (***Fig 5b***, left). The standard pipeline, however, exhibits less reproducibility; the five models in its ensemble are substantially less similar than GhostFold, particularly in the CDRH3 region, resulting in an ensemble CDRH3 RMSD of 4.79 Å (***Fig 5b***, right). This highlights a critical flaw in relying on sparse natural MSAs: the model produces a set of inconsistent, low-confidence hypotheses that are unreliable without extensive sampling and filtering. Indeed, even when we generated 20 models with the standard pipeline, only seven captured the correct CDRH3 conformation. GhostFold predicted near identical CDRH3 across all models.

This improvement in consistency is not an isolated case but a broader trend across the entire nanobody dataset (***Fig 5c***). We computed the sum of squares of RMSDs (ssRMSD) across the five-model ensemble for each target—a metric that penalizes both inaccuracy and inconsistency. GhostFold’s predictions (blue dots) consistently exhibit low ssRMSD values compared to AF2-MMseqs2 for long CDRH3 lengths and, forming a tight cluster near the bottom of the plot. In contrast, the predictions from the standard pipeline (red dots) show high variance and a clear trend of worsening performance as CDRH3 length increases. Even for targets with very long CDRH3 loops (>20), GhostFold predicted most of the loop right with dynamicity observed at the CDRH3 tip (9ISF, 9FJF and 9ENA) (***Fig 5d*** and ***Fig S5a–b***). For 9ISF (23AA CDRH3), GhostFold predicted the CDRH3 accurately, with deviation observed principally in CDRH2.

These findings demonstrate that by treating the CDRH3 loop as an “orphan peptide” and generating a structurally coherent pseudoMSA, GhostFold provides the necessary information for AlphaFold2 to bypass its reliance on extensive recycling and stochastic sampling. With a compact pseudoMSA of fewer than 100 synthetic sequences, we transform antibody modeling from a resource-intensive sampling problem into a highly accurate and deterministic prediction.

### GhostFold is computationally efficient

We next sought to evaluate several ways of further optimizing GhostFold’s performance and efficiency. We first tested the hypothesis that simply increasing computational effort would improve performance on difficult targets. Using our nanobody benchmark, we found that increasing the number of pseudoMSA generation blocks from one to five yielded no discernible improvement in CDRH3 prediction accuracy (***Fig 6a***). This suggests that the information content of the initial pseudoMSA is the critical determinant of success, and that brute-force increases in model depth offer diminishing returns.

**Figure 6.**
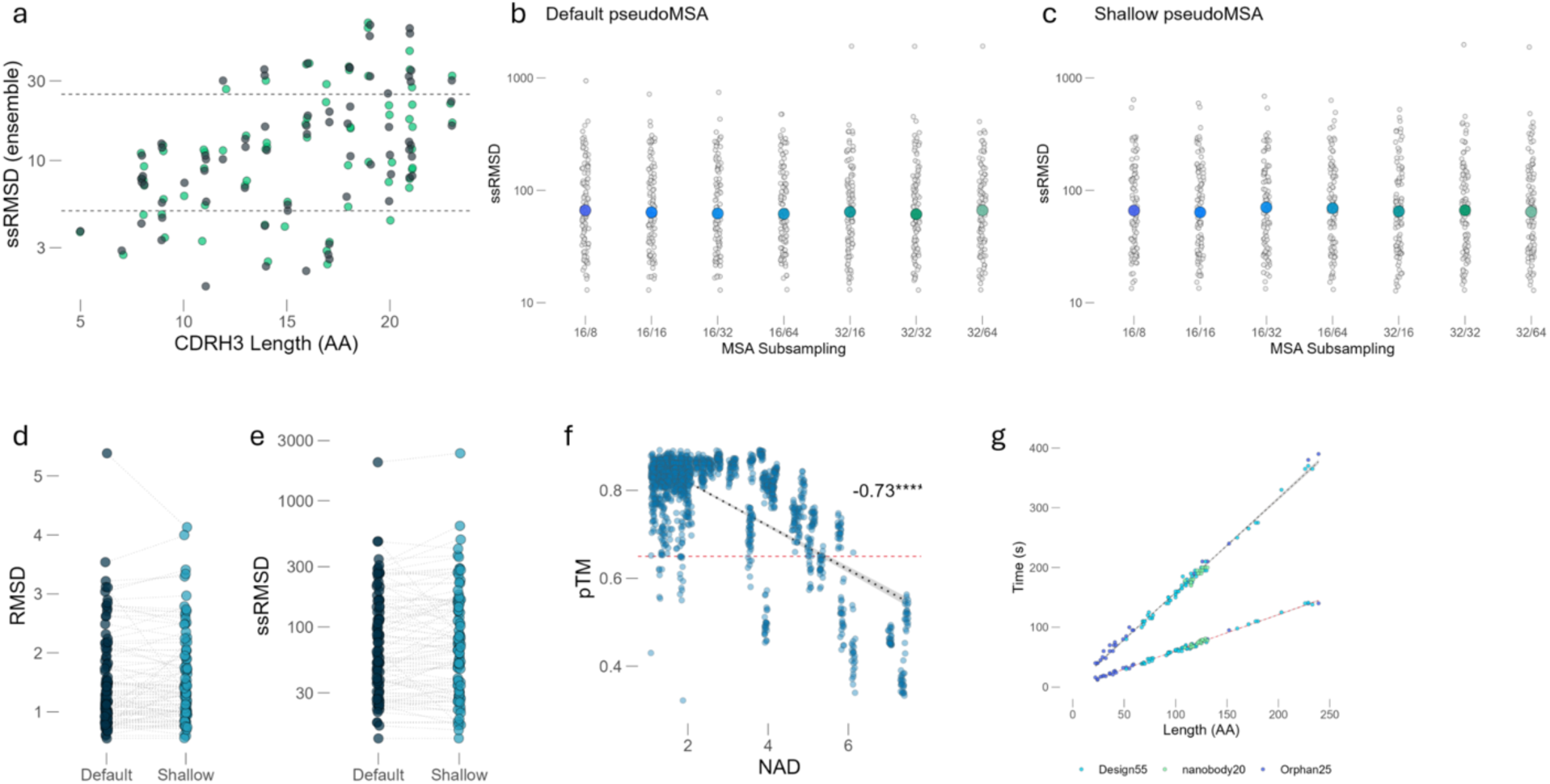
GhostFold enables compact, deterministic, and computationally efficient structure prediction. (a) Ensemble prediction variability (ssRMSD) of CDRH3 loops across the nanobody benchmark set, stratified by loop length. Increasing the number of pseudoMSA generation blocks (depth of structure conditioning) yields no improvement in prediction consistency, suggesting diminishing returns beyond a single block. Dashed lines indicate heuristic thresholds for low (5), moderate (25) ensemble variability. (b–c) Effect of subsampling depth on prediction consistency across the compiled100 dataset (comprised of Design55, Orphan, and Nanobody2025). Each dot represents the ensemble ssRMSD for a single target under a given sampling condition (max_seq/max_extra_seq), using the Default protocol (b) or the Shallow protocol (c). Colored dots represent median values for each group. Prediction quality remains stable across all tested sampling depths, even at minimal alignment sizes (e.g., 16/32), highlighting the information density of the synthetic pseudoMSA. (d–e) Direct comparison of prediction accuracy (RMSD) (d) and prediction consistency (ensemble ssRMSD) (e) between Default and Shallow pseudoMSA generation protocols. Each point corresponds to one target; lines connect the same target across protocols. RMSD and ssRMSD distributions are nearly identical, confirming that halving the number of generated sequences does not compromise performance. (f) Strong negative correlation between predicted model confidence (pTM) and the Normalized Alignment Divergence (NAD) score of the input pseudoMSA (Pearson R = –0.73, ****P < 1e–10). Lower NAD scores indicate greater pseudoMSA coherence and diversity balance, correlating with more confident structure predictions. The red dashed line indicates a pTM threshold of 0.7. (g) Total runtime for pseudoMSA generation as a function of input sequence length across the three benchmark sets. GhostFold shows linear scaling and completes MSA generation in under 400 seconds even for ∼250-residue targets (black line). The Shallow protocol (in red) achieves ∼3× speedup compared to the Default pipeline, enabling rapid structure prediction even for large proteins.

This led us to hypothesize that our generated pseudoMSAs are highly “distilled,” in the sense that a small number of sequences contain sufficient information for accurate prediction. To test this, we subsampled the pseudoMSAs for a comprehensive benchmark set (compiled100, combining orphan, design55, and nanobody2025) to varying depths (by varying the max_seq:max_extra_seq ratio where max_seq denotes the number of cluster representatives while max_extra_seq provide additional evolutionary couplings). Remarkably, even with just 16 sequences (and 32 extra sequences), the median prediction accuracy (RMSD) was nearly identical to that achieved with 32 sequences (and 64 extra sequences) (***Fig 6b***). These findings indicate that the synthetic coevolutionary signal is information-rich and it allows for significant reductions in computational overhead without sacrificing performance, confirming that the GhostFold pseudoMSA contains a highly potent and compact structural signal, making large alignments unnecessary.

Based on this insight, we engineered a more efficient “Shallow” protocol by reducing the number of sequences generated within the pseudoMSA block by half (from ∼450 to ∼225). This modification halved the VRAM requirement for the MSA generation step. When we repeated the subsampling experiment with this shallow protocol, we observed the exact same trend: performance remained stable even at very low subsampling depths (***Fig 6c***). A direct comparison of the final model accuracy for the “Default” and “Shallow” protocols showed virtually identical RMSD distributions (***Fig 6d***).

A key advantage of a strong input signal is prediction consistency. A common strategy to improve AlphaFold2 predictions is to generate a large ensemble of models (i.e., using different random seeds) and select the most representative structure. This, however, is computationally expensive. We assessed the convergence of a 25-model ensemble (five models for each of five random seeds) by calculating the sum of squares of RMSD (ssRMSD), penalizing both inaccuracy and inconsistency. For both the Default and Shallow protocols, the ssRMSD was extremely low indicating that all models in the ensemble converged to the same high-accuracy structure (***Fig 6e***), and nearly identical across two pseudoMSA generation protocols. This demonstrates that the GhostFold pseudoMSA provides a deterministic signal that obviates the need for extensive ensemble sampling, representing another significant computational saving.

To further ensure the reliability of our generative process, we sought to correlate the properties of the pseudoMSA with final model quality. We introduced a metric to rank pseudoMSAs called Normalized Alignment Divergence (NAD) score, which measures the consensus and diversity of the generated sequences, and serves as an effective quality filter (see ***Methods***). A lower NAD score, indicating a more converged and confident pseudoMSA, correlates well with higher final model confidence (pTM) (***Fig 6f***, R = -0.73). This allows for an upfront assessment of the input quality before committing to the full structure prediction pipeline.

Cumulatively, these optimizations result in a dramatic increase in computational efficiency. The “Shallow” protocol reduces the pseudoMSA generation time by nearly three-fold compared to our “Default” pipeline (***Fig 6g***). Furthermore, the pseudoMSA generation step is highly efficient, scaling linearly with protein length and typically completing in minutes, thereby removing the hours-long bottleneck associated with homology searching. For a typical 100-amino-acid monomer, the entire process takes approximately 55-65 seconds (0.55 – 0.65 seconds per residue). By generating a distilled, high-potency pseudoMSA, GhostFold transforms structure prediction from a resource-intensive sampling problem into a fast, deterministic, and highly accurate workflow.

## DISCUSSION

In this work, we introduce GhostFold and show that structure-constrained synthetic MSAs can effectively replace natural evolutionary data for accurate protein structure prediction. This is particularly useful when predicting the structures of orphan proteins or other classes of proteins for which sequence homologs are rare or non-existent. The prevailing paradigm has been to perpetually expand sequence databases, a strategy that demands massive computational resources (AlphaFold2’s standard databases are over 2 terabytes in size) and further increases yield diminishing returns. GhostFold represents a pivot away from this strategy of database expansion to one of structure-aware data generation, providing a robust, scalable, and computationally efficient solution to this long-standing challenge in a zero-shot fashion, requiring only a single query sequence.

The success of GhostFold is rooted in its novel use of a protein language model, ProstT5, to ‘backtrack’ evolution from a predicted latent structural scaffold. The initial projection of a sequence into its discrete structural alphabet captures the low-energy backbone conformation, and the subsequent inverse folding with ProstT5 generates a diverse ensemble of sequences predisposed to adopt this fold. The effectiveness of this approach is critically dependent on the design of the 3Di alphabet itself, which, unlike traditional structural alphabets that describe backbone torsion angles, encodes tertiary residue-residue interactions, albeit at a lower resolution. This design confers key advantages: it sharpens information density in conserved protein cores while minimizing it in variable loops, leading to a more potent signal for folding. The resulting pseudoMSA is not a random collection of mutations but an alignment rich with the emergent coevolutionary signals that the Evoformer architecture is trained to interpret, demonstrating that the essential information for folding is not evolutionary history *per se*, but the underlying structural constraints that coevolution encodes.

Our benchmarks reveal that optimizing the input data can fundamentally alter the behavior and efficiency of the prediction model. On challenging targets like the Design55 dataset, GhostFold’s pseudoMSAs provide such a complete signal that it renders AlphaFold2’s iterative recycling, a process critical for refining predictions from sparse inputs, largely redundant. This suggests that a significant portion of the model’s computational effort is a compensatory mechanism aimed at overcoming information-poor MSAs. By “frontloading” structural information into a high-quality synthetic MSA, we enable a more direct and efficient path to the final structure. This reframes the challenge of structure prediction, suggesting that future gains may come as much from input optimization as from architectural innovation. Encouragingly, GhostFold consistently matches or exceeds the performance of AlphaFold2 even on inputs for which deep and diverse MSAs can be built, meaning there is no performance trade-off for its significant gains in speed and accessibility.

A particularly important insight from our study is the robustness and generalizability of the GhostFold architecture. While several domain-specific structure prediction models, such as OpenFold, trRosettaX, RGN2, and others, have shown strong performance on targeted problems, they often do so at the cost of generality or extensibility. We demonstrate that our pseudoMSA-based approach not only outperforms single-sequence predictors like ESMFold and OmegaFold on orphan proteins, but also never underperforms these methods in any meaningful or consistent way. This universality makes GhostFold an attractive replacement for homology-based pipelines, while preserving compatibility with the broader AlphaFold2 framework. GhostFold either improves performance or holds steady; it rarely, if ever, does worse. This feature is particularly critical for downstream applications such as high-throughput prediction, protein design, and synthetic biology, where model generality is a core requirement.

One of the most surprising findings of this study is the observed decoupling of AlphaFold2’s pLDDT confidence metric from actual model accuracy when using our synthetic MSAs. We hypothesize that this decoupling arises because the confidence module has been implicitly trained to recognize the statistical patterns of natural evolutionary diversity. GhostFold’s alignments, while structurally informative, are “unnatural,” thereby confounding the confidence head. This insight may explain related phenomena, such as the low pLDDT scores often assigned to viral proteins, which evolve rapidly and lack deep MSAs, even when their predicted structures are correct [26]. This finding suggests caution against overreliance on pLDDT as a universal measure of prediction accuracy, particularly in contexts outside of natural protein families like de novo design and underscores the urgent need for new confidence metrics based more purely on geometric and stereochemical principles. Toward that end, we propose a new class of composite confidence metrics that integrate synthetic alignment quality with geometric indicators. Specifically, we suggest combining our Normalized Alignment Divergence (**NAD**) score, a proxy for pseudoMSA confidence and convergence, with geometric metrics such as ensemble RMSD variance, per-residue B-factor analogs, and low-energy clash scores. These hybrid metrics can be used to better assess prediction reliability in domains where pLDDT calibration fails. Future work could establish formal calibration schemes that jointly consider synthetic diversity, convergence, and structural determinism.

The practical utility of GhostFold is perhaps best demonstrated with antibody modeling—a notoriously difficult task due to the hypervariable and structurally dynamic nature of the CDRH3 loop. By treating the CDRH3 loop as an “orphan peptide” and generating a localized, structurally coherent pseudoMSA, GhostFold provides the necessary constraints for AlphaFold2 to model the loop with high accuracy. With a compact pseudoMSA of fewer than 100 synthetic sequences, we transform antibody modeling from a resource-intensive sampling problem into a highly accurate and deterministic prediction. This breakthrough effectively resolves a primary challenge that has historically hindered antibody-antigen docking and engineering. It is worth noting that AlphaFold2, in its single-sequence mode (i.e., without any MSA or coevolutionary signal), completely fails to model antibodies and nanobodies accurately, a limitation we have observed consistently in unpublished data. Future improvements to this approach may be achieved by further refining the NAD score specifically within complementarity-determining regions (**CDRs**), which exhibit highly localized structural diversity. By introducing position-aware or loop-sensitive adjustments to the NAD metric, it may be possible to improve pseudoMSA filtering and confidence assessment for hypervariable regions. This targeted refinement would allow even greater precision in synthetic alignment quality control, further enhancing accuracy and consistency in loop modeling.

Our “oracle” experiment, where near-perfect MSAs were generated from ground-truth structures using ProteinMPNN, places GhostFold’s performance in a remarkable context. A significant fraction of targets in the oracle set still failed to be predicted accurately, indicating that even a perfect representation of residue contacts does not guarantee a perfect prediction. While we assume that ProteinMPNN-generated sequences are structurally consistent with the target backbone, it is also possible, though less likely, that inaccuracies in the sequence design process introduced subtle deviations that undermined final model performance. Nonetheless, the fact that GhostFold, operating without any prior knowledge of the native structure, performs only marginally worse compared to this best-case scenario suggests our method is exceptionally effective at capturing the most critical information required for folding, and an inherent limit of predictability exists for certain topologies within the current AlphaFold2 framework. This points toward future research avenues aimed at improving the core folding algorithm itself, beyond the input MSA.

While prior work has shown that MSA augmentation with natural sequences improves prediction quality [12] or enhancing MSA quality by extensive database searching [2,3], our results demonstrate that similarly information-rich alignments can be synthetically generated. This raises an important practical point: improvements in protein structure prediction may be achieved not solely by deeper mining of evolutionary databases, but with biologically grounded synthetic data generation. Just as large language models in NLP have advanced dramatically through synthetic pretraining data and learned augmentation, we believe structure-aware generative frameworks should be actively explored and standardized. GhostFold is an early step in that direction, and future work should establish community guidelines, evaluation benchmarks, and composability criteria for synthetic protein alignments.

A central goal in developing new computational methods is not only to improve accuracy but also to enhance accessibility and efficiency. The entire workflow of shallow pseudoMSA with early stopping (most predictions converge within first recycle) in AlphaFold2 structure head significantly reduces the computational resources required for high-accuracy prediction and can be executed on a standard workstation equipped with a single consumer-grade GPU. The primary storage requirement is for the ProstT5 (∼12 GB) and AlphaFold2 (∼5.5 GB) model checkpoints, a reduction of several orders of magnitude compared to the conventional AlphaFold2 workflow that depends on multi-terabyte sequence databases. This makes GhostFold a much more lightweight and computationally accessible solution.

In conclusion, this work establishes structure-guided synthetic data as a powerful paradigm for addressing the limitations of homology-based structure prediction. GhostFold democratizes high-accuracy structural biology, making it accessible for any protein, regardless of its evolutionary history, and obviating the need for massive database installations. Looking forward, there is potential to further enhance the pseudoMSA generation by developing a richer structural tokenization alphabet designed explicitly for this task, rather than structure-based homology searches [16]. GhostFold paves the way for predictive models powered by strategic data generation, shifting the focus from large-scale collection and sheer volume to efficiency in advancing scientific discovery.

## LIMITATIONS

Despite GhostFold’s promising performance in structure prediction from synthetic MSAs, several limitations remain:

1. <u>Monomer-Only Support</u>: The current method supports only monomeric predictions, limiting applicability to multimeric assemblies and protein–protein interfaces. Multimer support is under development.
2. <u>No Template Integration</u>: GhostFold excludes structural templates, which may limit accuracy where homologous templates exist and improve predictions (TBM, template-based modelling category from CASP). Hybrid approaches could enhance future performance.
3. <u>Confidence Metric Discrepancies</u>: AlphaFold2’s internal confidence scores (pLDDT, pTM) are less predictive under synthetic alignments, complicating downstream validation. Alternative measures like NAD are promising but need further validation.
4. <u>Residual Structural Noise</u>: Although the pseudoMSAs are structurally informed, stochastic back-translation introduces noise, particularly in flexible or complex regions and refining or clipping that could enhance performance.
5. <u>Model and Alphabet Constraints</u>: GhostFold relies on ProstT5 and the 3Di alphabet, which were not explicitly optimized for fine-grained structure generation. Future iterations may benefit from purpose-built models and alphabets.
6. <u>Limited Modality Testing</u>: Generalization to disordered, membrane-associated, or modified proteins remains untested and may require tailored strategies.

These constraints highlight areas for future development, including multimer prediction, improved confidence calibration, and broader applicability across protein classes.

## METHODS

### Datasets

For this study, the Orphan and Design55 benchmarking datasets were adapted from the work of Wang et.al., 2022 and Wu et.al., 2022. The first, Orphan, is composed of 24 non-redundant “orphan” proteins from Wang et. al., 2022 (excluding 7A5P, non-continuous chain) and 12 “orphan” proteins from Wu et.al., 2022. These are naturally occurring proteins added to the PDB after May 2020 that have no identifiable sequence homologs in the UniRef50_2018_03 database. The second test set, Design55, includes 55 human-designed proteins. These proteins were collected from the PDB and filtered to exclude sequences with homologs in the training data or the UniRef50_2018_03 database. Design55 excludes very small (<50 residues), very large (>300 residues), or topologically simple structures.

### Curation of a nanobody benchmarking dataset

A comprehensive set of candidate structures was compiled from the Research Collaboratory for Structural Bioinformatics (RCSB) Protein Data Bank (PDB) as of July 2025. An initial query using the keyword “nanobody” was performed to retrieve all depositions potentially containing a VHH domain. To automate the isolation of nanobody chains from their cognate antigens or other complex members, we developed a heuristic-based computational pipeline.

Each retrieved PDB entry, defined as a structure S, was programmatically parsed using the BioPython library (v1.83) to iterate through its constituent polypeptide chains, {C_i_ ∣ i ∈ [1,n]}. For each chain C_i_, we defined its length L(C_i_) as the cardinality of the set of its standard amino acid residues, thereby excluding heteroatoms and solvent molecules. A chain was designated a candidate nanobody if its length satisfied the condition 80 ≤ L(C_i_) ≤ 140 residues. This length constraint was chosen based on empirical observations of VHH domain sizes.

To ensure the structural integrity of the candidates and exclude chains with unresolved backbone regions, we implemented a stringent contiguity filter such that the set of residue sequence identifiers for the standard residues in a candidate chain C_i_ were defined as an ordered set R_i_ = {s_1_,s_2_,…,s_k_}. The chain was retained for the final dataset if and only if it was fully contiguous, satisfying the condition s_j+1_ − s_j_ = 1 for all j ∈ [1,k − 1]. Chains failing this check were discarded. Each validated nanobody chain was saved into a new, single-chain PDB file. This process yielded a curated set of non-redundant, single-chain PDB files representing the ground-truth experimental structures, {E}. Finally, the amino acid sequence for each curated structure E was extracted from Cα coordinates. All extracted sequences were subsequently compiled into a single multi-FASTA file, which served as the unified input for all benchmarking experiments.

### Generative construction of a pseudo-MSA

To address the challenge of protein structure prediction for sequences with sparse or non-existent evolutionary records, we have developed a novel framework for the *de novo* generation of a pseudo-Multiple Sequence Alignment (pseudoMSA). The core of our methodology is a two-stage, fragment-based generative process that systematically explores the sequence space around the local and global structural features implied by the query. Our method circumvents the need for traditional homology searches by leveraging a large protein language model (pLM) to create a diverse ensemble of structurally plausible sequences conditioned on a single query protein. The foundation of our approach is ProstT5, which is based on the ProtT5-XL-U50 model. ProstT5 was fine-tuned to perform bi-directional translation between the 20 canonical amino acids (AA) and a 20-state structural alphabet (3Di). This 3Di vocabulary describes the tertiary interactions of each residue with its spatially closest neighbor, rather than its backbone torsion angles. This learned relationship between primary sequence and 3D structure is the engine for our generative pipeline.

The generation of the pseudoMSA for a given query sequence, S_q_, of length L begins with a fragment-based strategy designed to introduce localized diversity. We define a set of coverage fractions, C = {c_1_,c_2_,…,c_k_}, where c_i_ ∈ (0,1). For each coverage value c, a contiguous subsequence, or fragment s_f_, of length l = [c⋅L] is selected from a starting position chosen uniformly at random from the interval [0, L − l]. This fragment is the initial input for our two-stage generative process. In the first stage, termed **structural abstraction**, the amino acid fragment s_f_ is translated into its corresponding 3Di representation. We provide the model with the input prompt <AA2FOLD> s_f_ and generate a set of N distinct 3Di sequences, D = {d_1_,d_2_,…,d_N_}. Each d_i_ ∈ D represents a unique, plausible structural interpretation of the input fragment s_f_. The number of generated 3Di sequences, N, is a user-defined parameter (num_return_sequences). In the second stage, **conditional sequence diversification**, each of the N generated 3Di sequences is used as a condition to generate new amino acid sequences. For each d_i_ ∈ D, we prompt the model with <FOLD2AA> d_i_ to perform a back-translation. From each structural representation d_i_, we generate a set of M unique amino acid sequences, S_i_′={s_i_,1′,s_i_,2′,…,s_i_,M′}. The integer M, or the multiplier, dictates the sampling depth for each structural hypothesis, allowing for extensive exploration of sequences that can adopt a similar local fold. This two-stage process results in a total of *N×M* new amino acid fragments generated from the initial seed fragment s_f_.

To ensure the pseudoMSA is populated with a rich diversity of sequences, the generation process is performed across a matrix of stochastic decoding configurations. We define a hyperparameter space including sampling **temperature** (T), which modulates the probability distribution of the next token, **top-k sampling**, which restricts the sampling pool to the k most likely tokens, and **nucleus sampling (top-p)**, which restricts the pool to a cumulative probability mass p. The entire two-stage generation is iterated over the Cartesian product of these hyperparameter sets, creating a broad ensemble of sequences generated under varying degrees of randomness and constraint.

Following generation, each new fragment s_ij_′ is re-aligned into a full-length sequence of length L by padding the N- and C-terminal ends with gap characters (-) based on the fragment’s original starting position, effectively aligning the amino acids in a structurally aware alignment rather than conventional sequence-based alignment. This step is necessary as the back-translation introduces a sequence diversity that would lead to spurious alignment if sequence-aware aligners are used. A stringent filtering step is then applied to remove any generated sequences that do not perfectly match the target length L or have spurious and/or repetitive elements (high entropy), ensuring the integrity of the final MSA.

As an optional final step to introduce variation patterns that mimic natural evolution (randomized evolutionary drift), the filtered pseudoMSA can be further augmented. A user-defined percentage of the generated sequences are selected and subjected to mutation using standard substitution matrices, such as BLOSUM62, MEGABLAST and PAM250, at specified rates. This introduces evolutionarily informed substitutions into the generatively derived sequence set. The final pseudoMSA is constructed by concatenating the original query sequence S_q_ with the complete set of valid, full-length sequences generated across all coverage values, decoding configurations, and optional mutation steps from multiple independent runs. This comprehensive process yields a deep and diverse alignment, providing a robust input for downstream structure prediction.

### Beam ranking

To ensure a balanced evaluation that does not unfairly penalize longer, more complete sequences, the ranking of candidate sequences during beam search is determined by their length-normalized log-likelihood. The fundamental score for any given hypothesis is calculated as the sum of the log probabilities of its constituent tokens. However, because these log probabilities are inherently negative, this unnormalized sum creates a strong bias that favors shorter, often incomplete, sequences. To counteract this effect, we apply a length normalization penalty, which is achieved by dividing the cumulative log-likelihood of each sequence by its length raised to a penalty exponent, ɑ. This hyperparameter allows for fine-grained control over the degree of normalization, and for this work, it was tuned to ensure that sequences of varying lengths were evaluated equitably, thereby achieving a robust balance between probabilistic likelihood and sequence completeness.

### Structure Prediction Pipelines

Two distinct structure prediction pipelines were evaluated. The first, **ColabFold** (v1.5.5), was employed as a state-of-the-art baseline. The ColabFold pipeline utilizes MMseqs2 to rapidly generate a multiple sequence alignment (MSA) from a large sequence database, which is then used as input to the core AlphaFold2 model (alphafold_ptm) to predict the final structure. All predictions with ColabFold were executed with default parameters, with the following exceptions: six recycle steps, dropouts enabled, no templates To maintain consistency with the GhostFold protocol, post-prediction Amber relaxation was not performed.

The second pipeline, our **GhostFold** method, fundamentally departs from the homology-based approach. GhostFold utilizes the generated pseudoMSA (a synthetic alignment engineered to encapsulate the implicit co-evolutionary and physicochemical constraints of a sequence) without relying on a database search for extant homologs. This pseudoMSA is formatted to be compatible with the AlphaFold2 architecture and provided as the primary input feature, replacing the conventional MSA. This allows for structure prediction in the absence of a sufficiently deep MSA, a common challenge for proteins like orphan proteins and antibodies/nanobodies which may have sparse evolutionary histories.

### Structural comparison

The accuracy of our predicted protein models was evaluated by comparing them against their corresponding experimental ground truth structures from the RCSB PDB. We calculated the Root Mean Square Deviation (RMSD) between the backbone Carbon-alpha (Cα) atoms of the two structures. To ensure a fair comparison, even when the predicted and experimental structures had different numbers of residues, we first identified the set of Cα atoms common to both. The predicted structure was then structurally aligned to the ground truth structure by superimposing this common set of atoms and the final RMSD value was calculated for these aligned atom pairs.

### Quantitative assessment of ensemble convergence

To evaluate the performance of our pseudoMSA augmented pipeline, a quantitative analysis was performed to assess how well each prediction ensemble converged towards its respective ground truth structure. The primary metric for this assessment was the sum of squared Root-Mean-Square Deviations (RMSD) across all predictions in an ensemble. This metric provides a single, aggregate score that reflects both the accuracy (proximity to the native state) and the precision (spread of the decoys) of the prediction set. A lower score indicates better convergence on a structure close to the native state. The analysis was carried out using a custom Python script leveraging the Bio.PDB module from the Biopython library (version 1.83). The procedure for a single ensemble was as follows:

1. **Structure Parsing and Atom Selection:** Both the reference structure and each predicted decoy structure were parsed from their PDB files. To focus the comparison on the overall backbone fold and minimize the influence of side-chain packing variations, only the coordinates of the C-alpha (Cα) atoms were considered for the analysis.
2. **Structural Superposition and RMSD Calculation:** For each predicted structure, its Cα atoms were structurally aligned to the Cα atoms of the ground truth reference structure. This superposition was achieved using a least-squares fitting algorithm, as implemented in Biopython’s Superimposer class. Following the optimal rotation and translation, the RMSD was calculated by taking the square root of the mean of the squared distances between each pair of corresponding atoms.
3. **Ensemble Convergence Score:** To obtain the final convergence score for the ensemble, the individual RMSD value for each successfully compared decoy was first squared. These squared RMSD values were then summed together to yield the final score for the entire ensemble (ssRMSD). Squaring the RMSD gives greater weight to predictions that deviate more significantly from the native structure, thus effectively penalizing poorly converged ensembles.

### Normalized alignment divergence

To quantitatively assess the quality and consensus of the pseudo-multiple sequence alignments (MSAs) generated by our fragment-based methodology, we developed a metric we term Normalized Alignment Divergence (NAD). The generative nature of our approach, while powerful, can occasionally result in low-quality outputs if the ProstT5 back-translation model falls into a repetitive or non-convergent sampling mode. While a preliminary filter removes simple mono- or di-peptide repeats, the NAD score provides a more holistic measure of the alignment’s integrity. The calculation of the NAD score for a given MSA is performed as follows:

1. **Pairwise Sequence Identity Assessment**: The process begins with an all-vs-all comparison of the sequences within the alignment. For each pair of sequences, we calculated their fractional identity, defined as the number of positions with identical amino acid residues divided by the total length of the alignment (L).
2. **Homology Thresholding**: A sequence identity threshold was set at 0.5. Any pair of sequences with an identity score greater than or equal to this threshold was classified as homologous. This step effectively groups similar sequences into clusters.
3. **Individual Sequence Weighting**: A unique weight is assigned to each sequence to penalize redundancy. For each sequence i, we counted the number of other sequences homologous to it. The weight for sequence i, denoted w_i_, was then computed as the reciprocal of one plus this count. This method assigns a low weight to sequences in dense, homologous clusters and a weight approaching 1.0 to sequences that are distinct from others.
4. **Aggregation and Length Normalization**: The final NAD score is derived by summing the individual weights (w_i_) of all sequences in the alignment. To ensure scores are comparable across alignments of different lengths, this sum is normalized by multiplying it by the inverse of the square root of the alignment length (L).

The resulting NAD score provides a single, continuous value representing the alignment’s divergence. In the context of our generative pipeline, a low NAD score is desirable. It signifies a convergent, high-quality pseudo-MSA, indicating that the model is confident in its output because the generated sequences belong to a tight embedding cluster of a specific, well-defined family of sequences corresponding to the input 3Di representation. In contrast, a high NAD score signifies a divergent, low-quality alignment, potentially resulting from the model falling into a repetitive or low-quality sequence generation mode, suggesting the model is sampling broadly and is less certain about the optimal sequence. This high divergence implies that the sequences are scattered throughout the embedding space, suggesting the model is guessing the best sequence to fit the structural representation and reflecting low confidence in the prediction.

## CODE AVAILABILITY

GhostFold is MIT licensed and freely available at github.com/brineylab/ghostfold.

## AUTHOR CONTRIBUTIONS

NM and BB conceptualized the study. Data generation and analysis was performed by NM. The GhostFold software package was developed by NM. The manuscript was prepared, revised and reviewed by all authors.

---

**Supplementary Figure 1.**
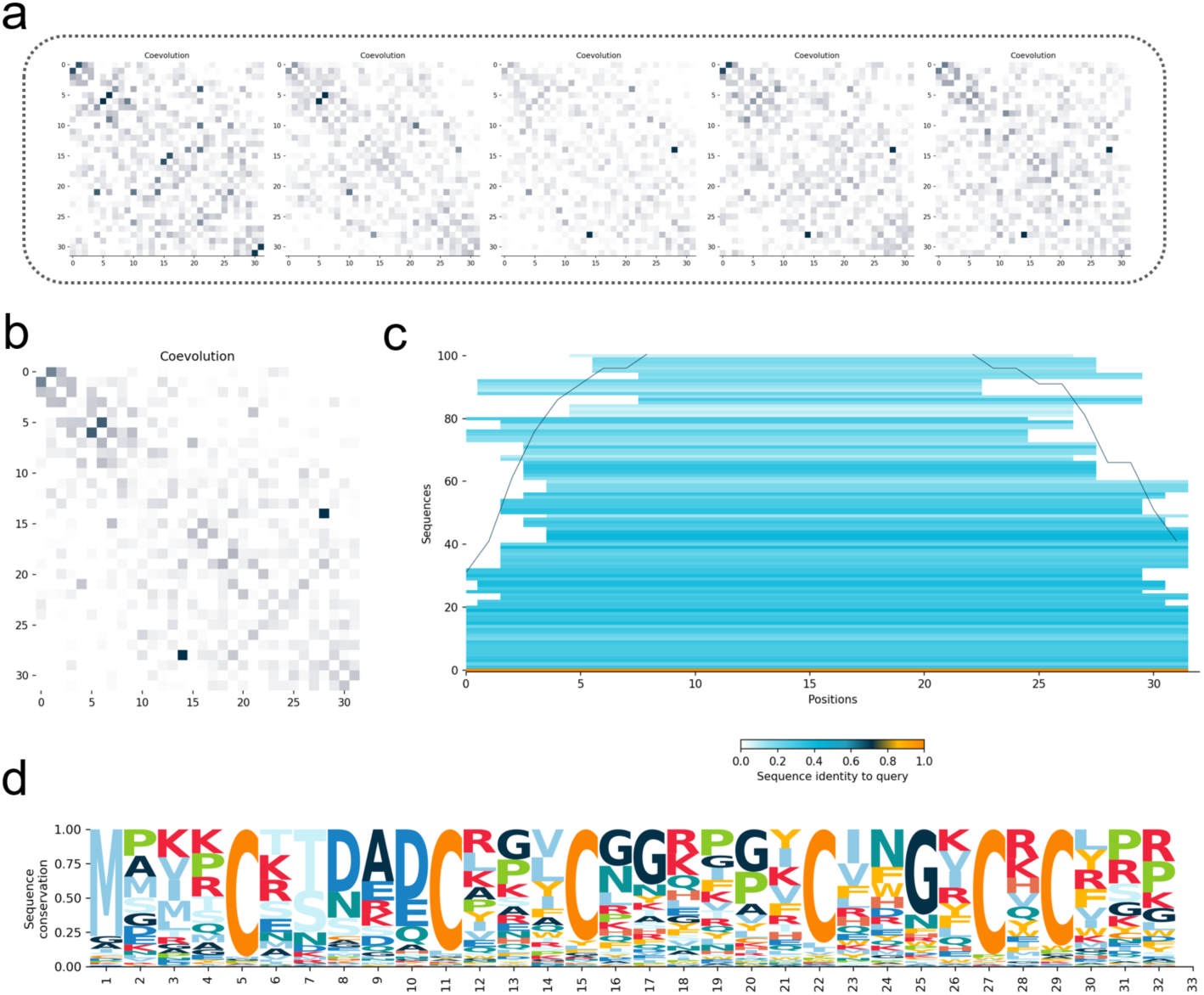
Emergence of coevolutionary signal and structural coherence in synthetically generated pseudoMSAs. (a) Pairwise contact probability matrices from representative back-translated sequences generated using the AA→3Di→AA pipeline, visualized as individual covariation heatmaps. Each map is sparse and noisy in isolation, capturing only weak coevolutionary signals due to the limited sequence length and stochastic nature of individual samples. (b) Aggregated coevolution matrix derived from the full pseudoMSA, constructed from 100 synthetically generated sequences. Despite originating from de novo generation, the alignment exhibits robust long-range coevolutionary couplings between distal residues—hallmarks of structural constraints encoded within the β-grasp fold of the input query (6MRQ). (c) Positional coverage and diversity of the generated pseudoMSA. Horizontal bars represent individual sequences, colored by sequence identity to the original query. Sequences span from full-length to partial fragments (coverage 40–100%) to enhance local structural motif diversity. The grey line indicates total coverage per position, showing uniform representation across the full sequence. (d) Sequence logo representing positional amino acid preferences across the pseudoMSA. The alignment captures conserved structural features of the β-grasp fold, including glycine and proline residues at key positions and terminal cysteines involved in disulfide bridge formation. The variability in other positions reflects controlled evolutionary drift induced via stochastic decoding and substitution-matrix-guided mutagenesis.

**Supplementary Figure 2.**
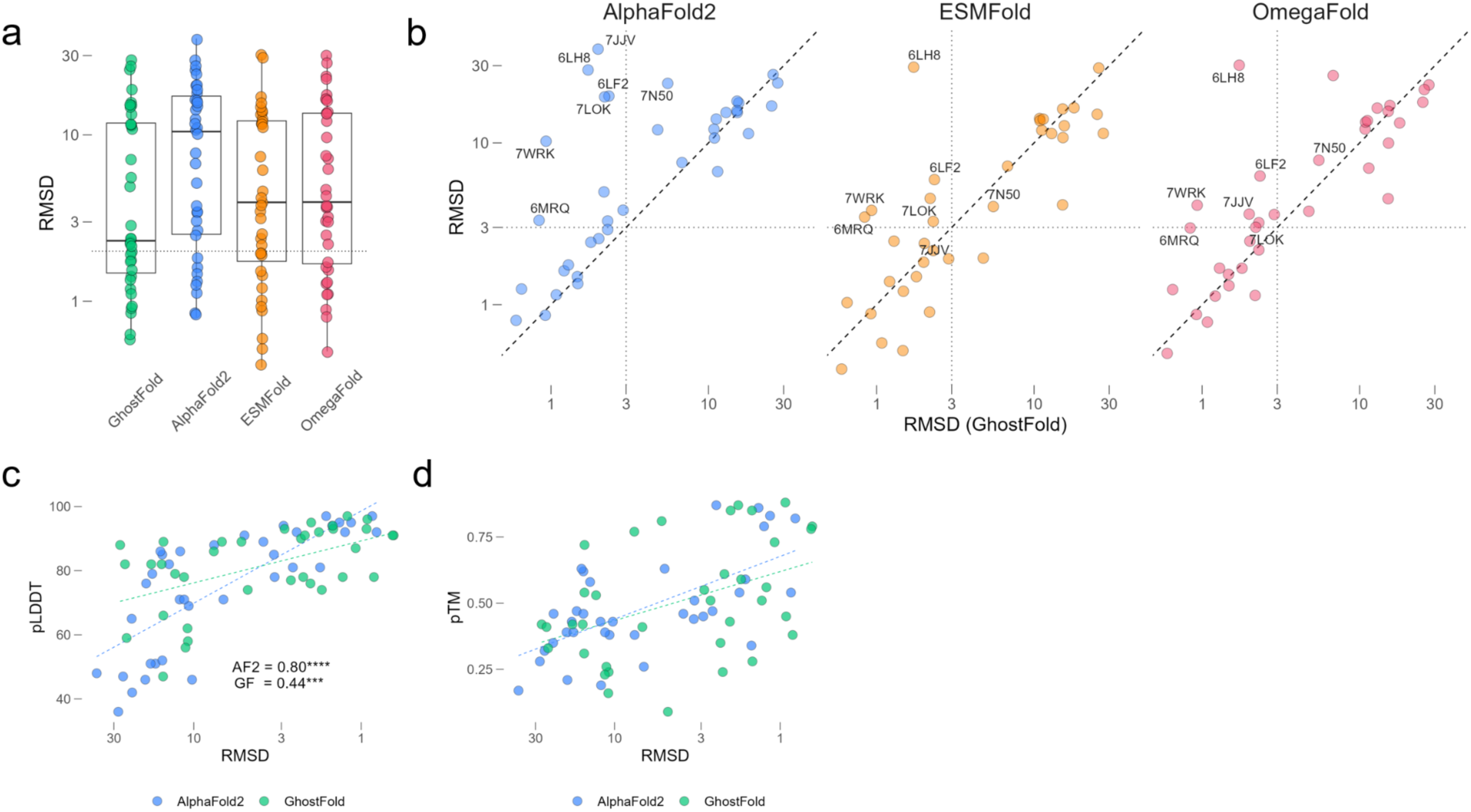
GhostFold yields consistently lower RMSD and alters AlphaFold2 confidence score interpretation. (a) Distribution of RMSD values for all 36 orphan protein targets across GhostFold, AlphaFold2, ESMFold, and OmegaFold. Boxplots show median, interquartile range, and individual data points (log scale). GhostFold displays a tighter and lower RMSD distribution, indicative of overall higher accuracy. (b) Extended RMSD comparison plots for GhostFold versus AlphaFold2, ESMFold, and OmegaFold, including all 36 orphan proteins. This includes the 12 “failure” cases where all predictors had RMSD > 3 Å. (c) Correlation between RMSD and per-residue confidence scores (pLDDT) for AlphaFold2 (blue) and GhostFold (green). While AF2 shows a strong negative correlation (R² = 0.80), GhostFold’s correlation, though significant, is noticeably weaker (R² = 0.44), indicating a potential shift in the interpretability of pLDDT scores when using synthetic pseudoMSAs. (d) Correlation between RMSD and predicted TM-score (pTM) for AlphaFold2 and GhostFold. GhostFold models tend to have lower pTM values overall (< 0.5), but a weak linear trend remains. The divergence in confidence metrics underscores the need for alternative confidence calibration when using pseudoMSAs.

**Supplementary Figure 3.**
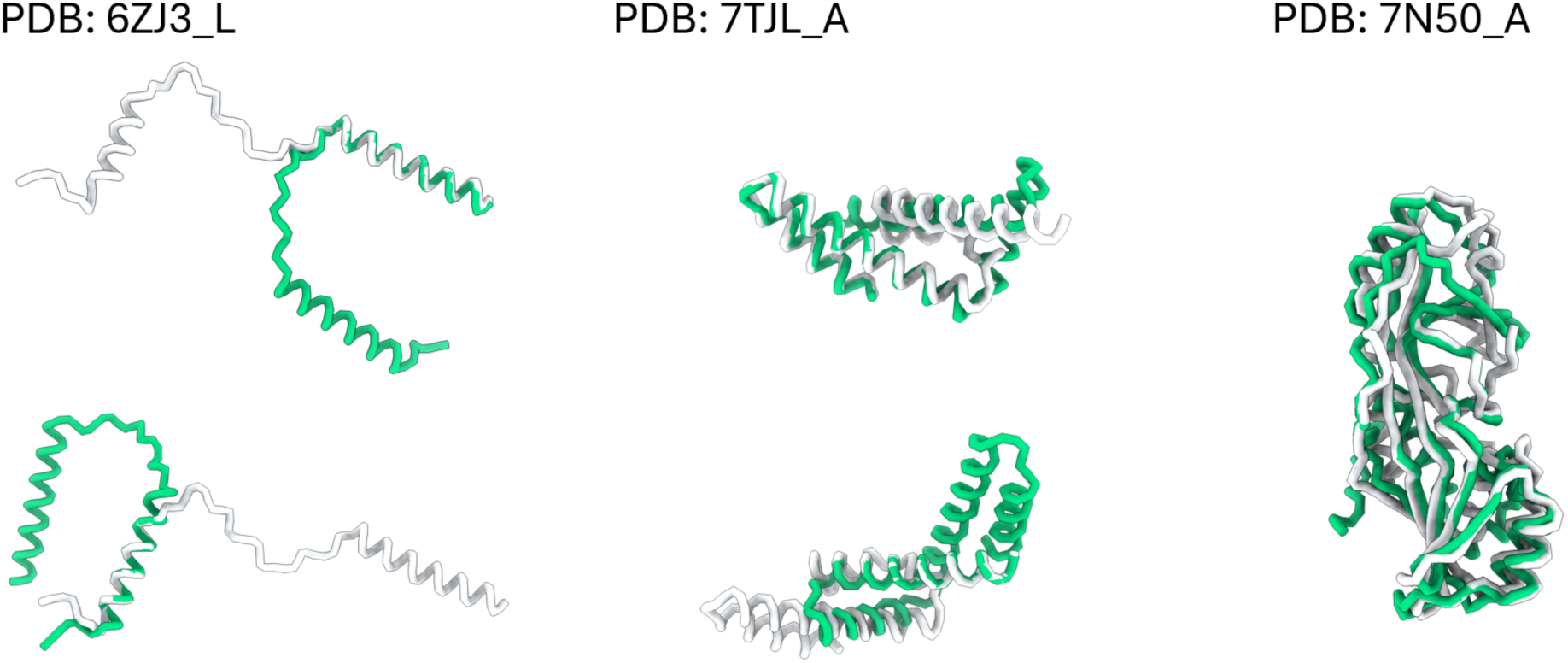
GhostFold recovers correct domains in globally “failed” targets. For 12 of 36 orphan proteins, no method produced a fully correct full-length model; nevertheless, GhostFold recapitulates most constituent domains in several of these failures. Shown are three representative cases (6ZJ3_L, 7TJL_A, 7N50_A). For each protein, shown are the global superposition of the native structure (gray) and the GhostFold prediction (green), illustrating incorrect inter-domain packing/relative orientation that inflates the full-chain RMSD.

**Supplementary Figure 4.**
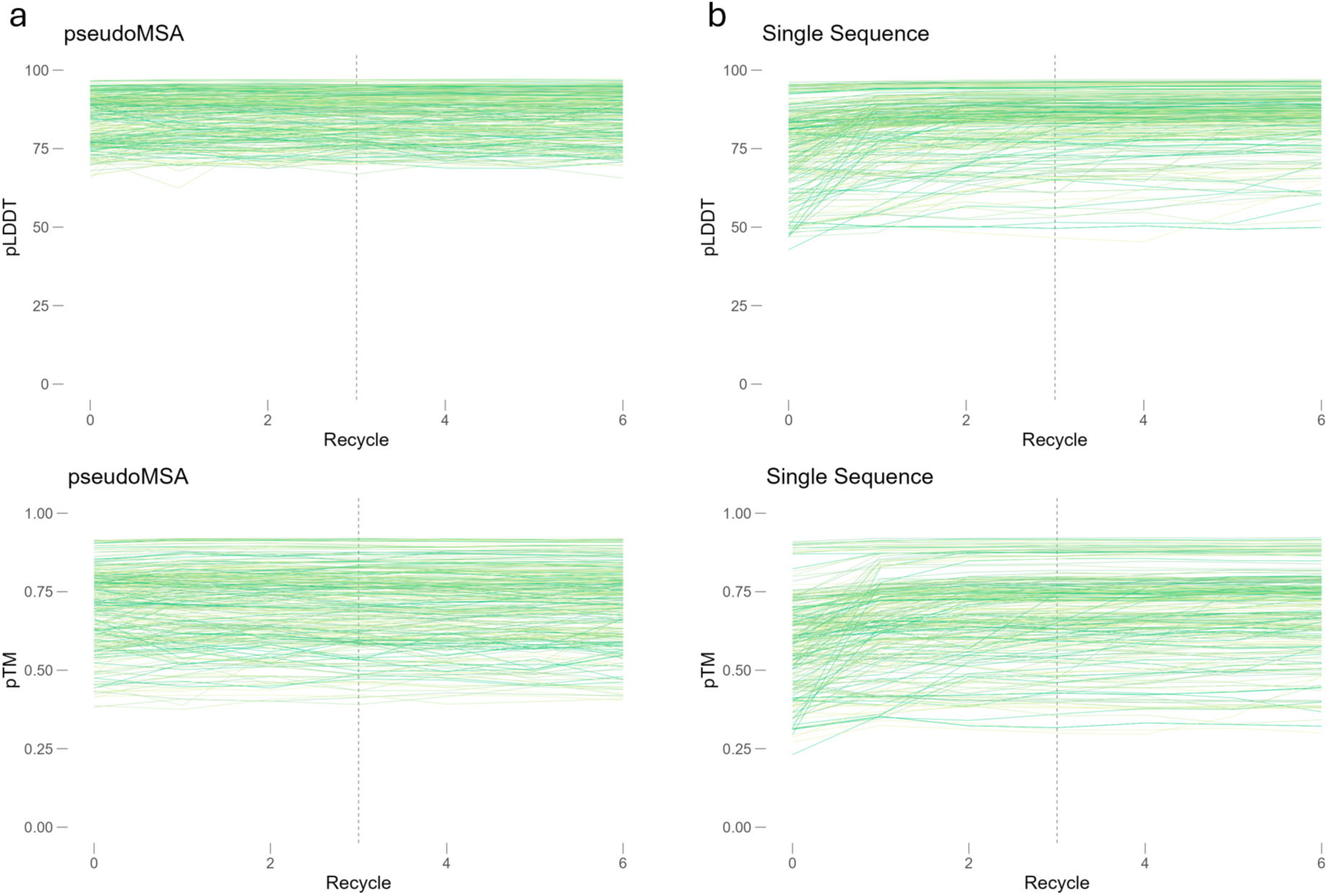
GhostFold yields consistently high confidence scores across all AlphaFold2 model variants. (a–b) Line plots showing pLDDT (top) and pTM (bottom) values across recycling steps (x-axis) for all five AlphaFold2 model variants (different shades of green) on the Design55 dataset. Each line represents one design prediction from a given model. (a) GhostFold (pseudoMSA input) maintains consistently high per-residue (pLDDT) and global fold (pTM) confidence across all five AlphaFold2 models. Confidence scores remain stable from recycle 0 onward, indicating minimal dependence on iterative refinement. (b) In contrast, the standard single-sequence mode (without pseudoMSA) exhibits greater variance and a broader dynamic range in both pLDDT and pTM values. Confidence scores generally increase with recycling, indicating that the model relies on iterative refinement to achieve convergence in the absence of strong input signal. Vertical dashed lines indicate the transition from early to later recycling iterations (recycle 3). Across all five model variants, GhostFold provides structurally informative pseudoMSAs that result in confident predictions from the very first recycle, underscoring the robustness of the approach across AlphaFold2’s model ensemble.

**Supplementary Figure 5.**
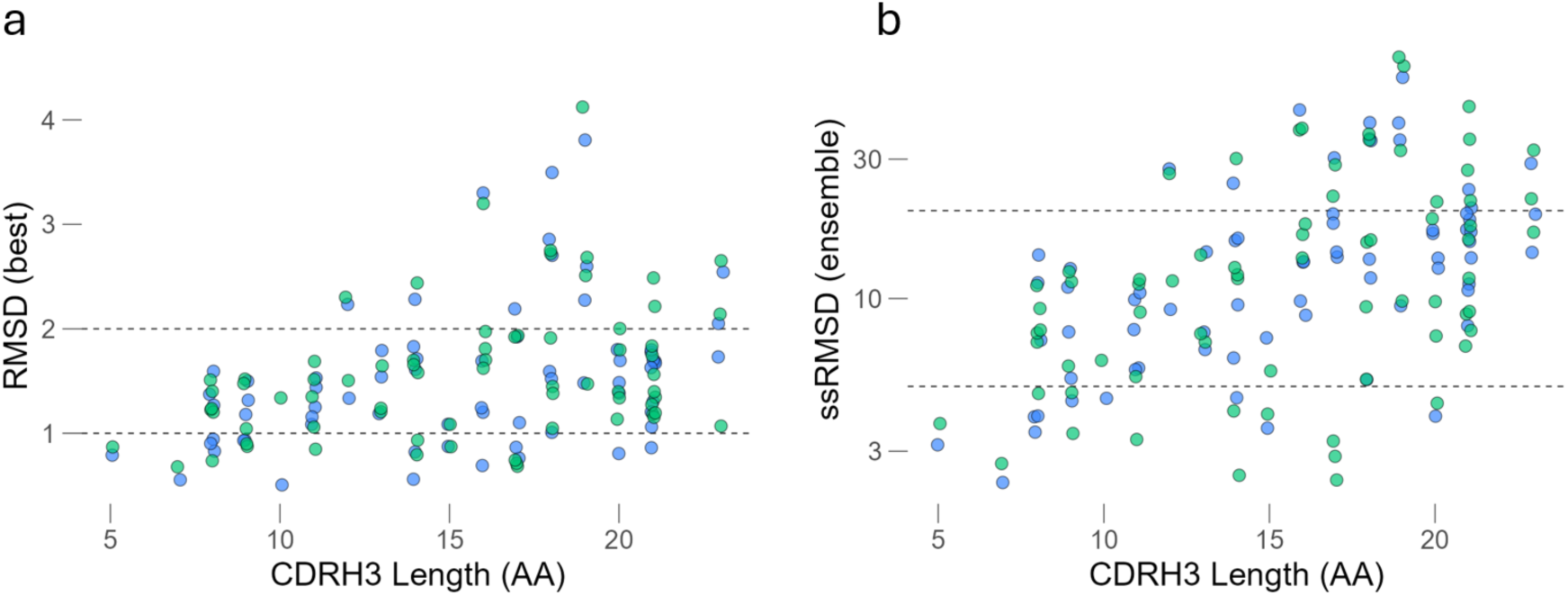
GhostFold ensemble consistency is robust to loop length and model variance. (a) Per-target best CDRH3 RMSD as a function of loop length. GhostFold (green) maintains low RMSD across increasing CDRH3 lengths, while AlphaFold2 (blue) shows degraded performance for loops longer than 15 residues. Dashed lines at 1 Å and 2 Å indicate accuracy benchmarks for loop modeling. (b) Ensemble ssRMSD plotted against CDRH3 loop length. GhostFold remains stable and compact across all lengths, while AlphaFold2 shows increasing variability with longer loops. Together with panel B, this highlights GhostFold’s unique ability to maintain both accuracy and consistency across difficult hypervariable regions.

